# Differential Expression of *NPAS4* in the Dorsolateral Prefrontal Cortex Following Acute Opioid Intoxication

**DOI:** 10.1101/2020.12.23.424239

**Authors:** David W. Sosnowski, Andrew E. Jaffe, Ran Tao, Amy Deep-Soboslay, Joel E. Kleinman, Thomas M. Hyde, Chang Shu, Sarven Sabunciyan, Brion S. Maher

**Author notes:** Corresponding Author: David W. Sosnowski, Ph.D., 624 N. Broadway, Hampton House, Baltimore, MD 21205, Phone: N/A.

## Abstract

**Background and Aims:** The physical, emotional, and social impacts of opioid abuse are well known; although preclinical models reveal the neurobiological pathways altered through opioid abuse, comprehensive assessments of gene expression in human brain samples are lacking. The goals of the present study were to compare gene expression in the prefrontal cortex between brain samples of individuals who died of acute opioid intoxication and group-matched controls, and to test if differential gene expression was enriched in gene sets related to opioid use.

**Design:** Cross-sectional study using human brains donated to the Lieber Institute for Brain Development. Study groups included 72 brain samples from individuals who died of acute opioid intoxication, 53 group-matched psychiatric control samples, and 28 group-matched normal control samples.

**Setting:** Maryland, USA.

**Participants:** Postmortem tissue samples of the dorsolateral prefrontal cortex from 153 deceased individuals *(Mage* = 35.42, *SD* = 9.43 years; 62% male; 77% White).

**Measurements:** Whole transcriptome RNA sequencing was used to generate exon counts, and differential expression was tested using *limma-voom.* Analyses controlled for relevant sociodemographic characteristics, technical covariates, and cryptic relatedness and batch effects using quality surrogate variable analysis. Gene set enrichment analyses (GSEA) also were conducted.

**Findings:** Sixteen genes were differentially expressed (i.e., FDR-corrected *p* < .10) in opioid samples compared to control samples. The top differentially expressed gene, *NPAS4* (FDR adjusted *p* = .005), was downregulated in opioid samples and has previously been implicated in cocaine use. Enrichment analyses did not provide evidence for enrichment in pathways obviously related to opioid use.

**Conclusions:** *NPAS4* is differentially expressed in the prefrontal cortex of subjects that died of an opioid overdose, providing evidence for another gene with functional relevance to opioid use and overdose.

## Introduction

According to the Centers for Disease Control and Prevention (CDC), more than 80% of drug overdose deaths involve opioids (1). Moreover, the use of illicitly manufactured fentanyl is rising, with two-thirds of opioid-related overdose deaths involving synthetic opioids (2). In addition to the loss of life, opioid abuse is estimated to cost society as much as $78.5 billion per year (3). The individual and societal costs of opioid abuse underscore the need to clarify the molecular underpinnings of opioid misuse, which will aid in the development of therapeutic targets and subsequent interventions to effectively treat opioid use disorder (OUD). Although preclinical models are an essential first step in understanding the neurobiological mechanisms underlying opioid abuse, examination of gene expression in human brain tissue is critical to accurately characterize neurobiological differences between opioid users and non-users (4). The goal of the present study was to compare genome-wide patterns of gene expression in the dorsolateral prefrontal cortex (dlPFC) of postmortem human adult brains among individuals who died of acute opioid intoxication and group-matched controls. The dlPFC was selected based on preclinical and clinical evidence demonstrating its role in addiction (5–7).

Research examining the neurobiology of opioid abuse has focused primarily on dopaminergic and glutamatergic neurotransmission within the nucleus accumbens (NAc) and prefrontal cortex (PFC) (7,8). Similar to other drugs of abuse, opioids exert their initial influence on addiction through a surge in dopamine within the NAc (5). Specifically, opioids activate μ-opioid receptors, disinhibiting neuron firing and leading to substantial increases in dopamine neurotransmission within the NAc (9). Prolonged use of opioids is also associated with similar changes in glutamatergic transmission (7); these changes are associated with cue reactivity (i.e., cravings) (10), deficits in learning, memory, and attention (11), and increased impulsivity, which can last for years after abstinence (12). Moreover, evidence demonstrates that the PFC and NAc interact in a manner such that dopamine-related adaptations in the PFC influence behavioral responses to drug-related stimuli, while simultaneous glutamate-related adaptations in the NAc contribute to compulsive drug-seeking behavior (6). These findings underscore the interdependence of these brains regions and value in exploring changes in gene expression and function within the PFC as a result of opioid abuse.

Although the molecular pathways and neurocognitive consequences of opioid abuse are well studied, changes in gene expression driving these adaptations are less well characterized. To date, only one study has examined genome-wide differences in gene expression in the human brain between opioid users and controls (13). The authors found 545 differentially expressed genes (FDR adjusted *p* < .10) in the midbrains of deceased opioid users, and the majority of these genes were protein coding genes that were upregulated. The study was limited, however, in that analyses did not adjust for cell composition or comorbid psychiatric diagnoses. The current study builds upon prior work in two ways. First, this analysis focuses on gene expression in the dlPFC. Although gene expression has been studied in the midbrain (13), and both the midbrain and dlPFC are implicated in addiction, evidence suggests that gene expression varies across brain regions (14). Second, the present analysis contains data on cell composition and comorbid psychiatric diagnosis, accounting for primary sources of confounding. Results will clarify patterns of gene expression in the PFC as a result of opioid overdose, informing subsequent efforts to identify therapeutic targets.

## Methods

### Participants and Procedure

Postmortem brain samples were donated to the Lieber Institute for Brain Development from the Offices of the Chief Medical Examiner of the State of Maryland (MDH protocol #12-24) and of Western Michigan University Homer Stryker School of Medicine, Department of Pathology (WIRB protocol #1126332); One brain sample was acquired through a material transfer agreement from the National Institute of Mental Health (NIMH; donated through the Office of the Chief Medical Examiner of the District of Columbia: protocol NIMH#90-M-0142), all with the informed consent of legal next-of-kin at the time of autopsy. The present study contained 160 postmortem human brain samples *(M_age_* = 35.15, *SD* = 9.42 years; 62% male; 78% White). Tables 1 and 2 provide detailed information on the study samples. Five psychiatric controls were removed for testing positive for an opioid at time of death, and one opioid sample and an additional psychiatric control were removed due to mismatching observed and predicted sex. The analytic sample contained 153 samples, consisting of 72 opioid-positive samples, 53 group-matched psychiatric controls, and 28 group-matched controls.

**Table 1.**
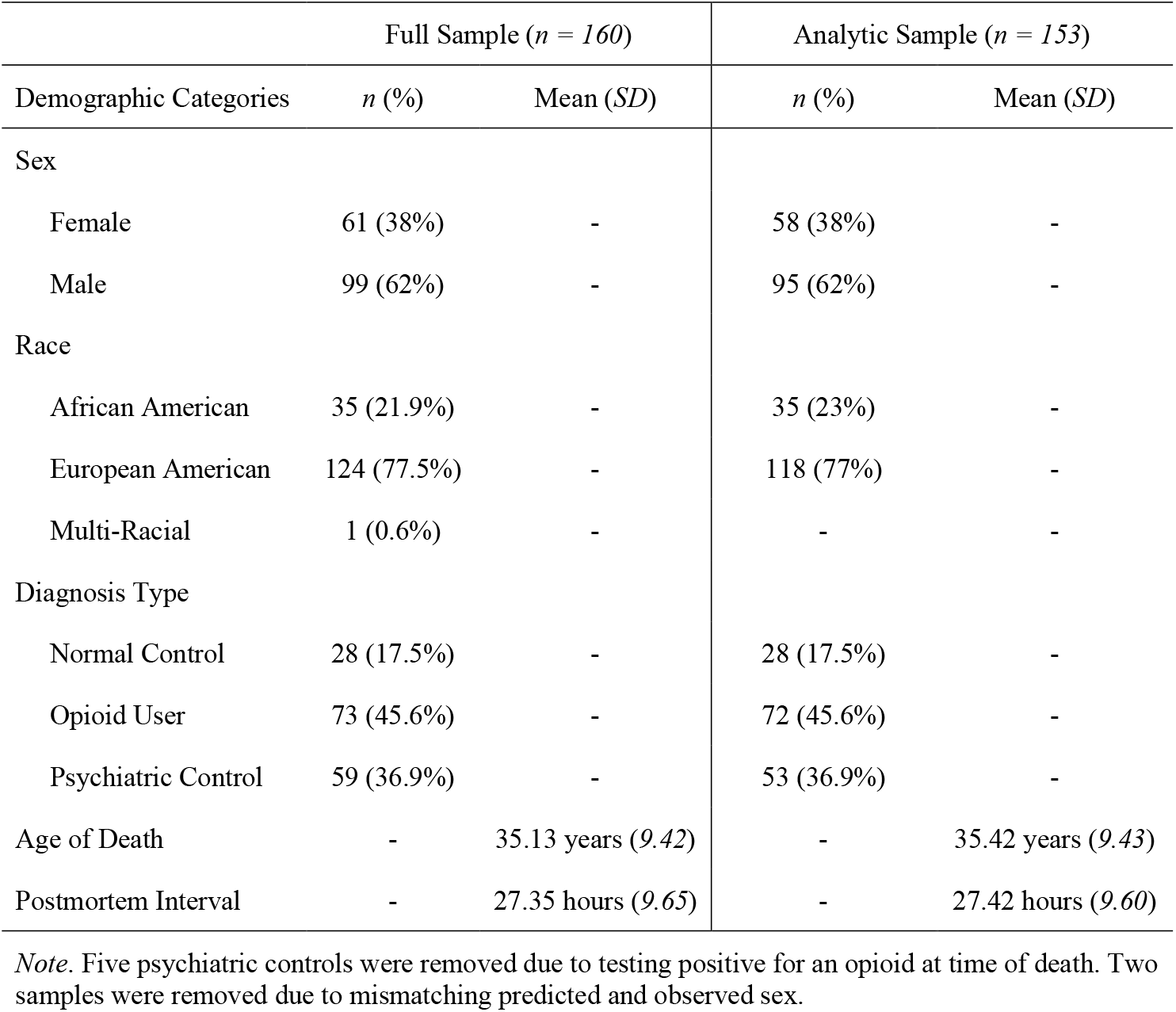
Sample Demographic Information

**Table 2.**
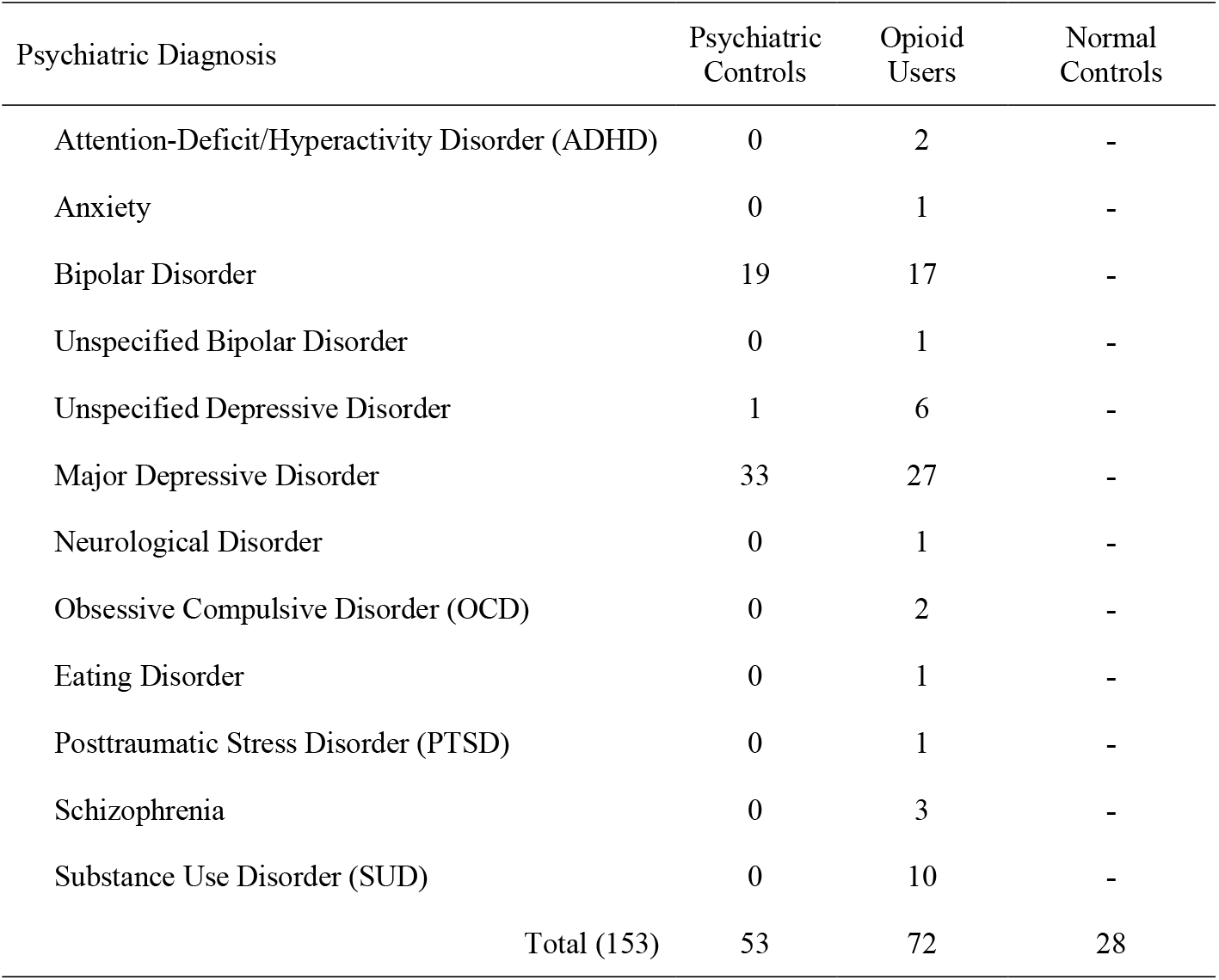
Primary Psychiatric Diagnosis by Study Group (n = 153)

At the time of donation, a 36-item next-of-kin informant telephone screening was conducted to obtain medical, social, demographic, and psychiatric history. Macroscopic and microscopic neuropathological examinations were conducted on every case by a board-certified neuropathologist to exclude for neurological problems, neuritic pathology, or cerebrovascular accidents. A retrospective clinical diagnostic review was conducted on every brain donor, consisting of the telephone screening, macroscopic and microscopic neuropathological examinations, autopsy and forensic investigative data, forensic toxicology data, extensive psychiatric treatment, substance abuse treatment, and/or medical record reviews, and whenever possible, family informant interviews.

All data were compiled into a comprehensive psychiatric narrative summary that was reviewed by two board-certified psychiatrists in order to arrive at lifetime DSM-V psychiatric diagnoses (including substance use disorders/intoxication) and medical diagnoses. Nonpsychiatric healthy controls were free from psychiatric and substance use diagnoses, and their toxicological data were negative for drugs of abuse. Every brain donor had forensic toxicological analysis, which typically covered ethanol and volatiles, opiates, cocaine/metabolites, amphetamines, and benzodiazepines. Some donors also received supplemental directed toxicological analysis using National Medical Services, Inc., including nicotine/cotinine testing, cannabis testing, and the expanded forensic panel in postmortem blood (or, in rare cases, in postmortem cerebellar tissue) in order to cover any substances not tested. The following substances were considered opioids: codeine, morphine, oxycodone, hydrocodone, oxymorphone, hydromorphone, methadone, fentanyl, 6-monoacetylmorphine, and tramadol.

### RNA Measurement and Preprocessing

RNA was concurrently extracted with DNA from 100mg of dlPFC tissue using the QIAGEN AllPrep DNA/RNA Mini Kit. Paired-end strand-specific sequencing libraries were prepared from 300ng total RNA using the TruSeq Stranded Total RNA Library Preparation kit with Ribo-Zero Gold ribosomal and mitochondrial RNA depletion. Resulting RNA-seq libraries were sequenced on an Illumina HiSeq 3000 at the Lieber Institute for Brain Development Sequencing Core. Raw sequencing reads were processed into gene counts using a previously described pipeline (15). Briefly, reads were aligned to the genome using HISAT2 (16) and the 58,037 Gencode v25 genes were quantified from resulting alignments using featureCounts using paired-end stranded counting (17). We calculated the reads per kilobase of transcript, per million mapped reads (RPKM) from these counts, and genes with an average RPKM < .20 were excluded from analysis. This resulted in 25,644 genes used in the differential expression analysis. We then calculated principal components (PCs) and examined correlations between the PCs and phenotypic and technical covariates. The first two principal components, accounting 25.7% of variance in the data, were significantly correlated with all technical covariates (e.g., overall mapping rate, RNA integrity number [RIN]), and the first PC was significant correlated with observed sex. Thus, these variables were included among the list of covariates in the differential expression analysis.

### Statistical Methods

#### Differential expression

Group-wise differences in gene expression between opioid-positive samples and controls were tested in R version 3.6.1 (18) using *limma-voom* (19). Specifically, a linear model was fit where gene expression was regressed on opioid use status (yes/no) across each of 25,644 expressed genes. The model adjusted for the following covariates: diagnosis of a psychiatric disorder other than a substance use disorder (yes/no), age of death, sex, postmortem interval (PMI), cell composition (i.e., % positive neurons), sex, race (Black vs. White), seven technical covariates (e.g., RIN), and 10 surrogate variables identified through quality surrogate variable analysis (qSVA) (20). Analyses were corrected for multiple testing and statistical significance was determined by an FDR corrected *p*-value < .10 (21).

#### Gene set enrichment analysis

Gene set enrichment analysis (GSEA) was conducted to explore in which biological pathways, or gene sets, enrichment of differential gene expression existed. Following procedures outlined by Reimand et al. (22), we extracted differential expression results (i.e., gene symbols, log fold-change [logFC], nominal *p*-values) comparing opioid and control samples and created a ranked list of all genes included in our experiment (*n* = 25,644). We ranked genes by first obtaining values of 1, 0, and −1 representing logFC values above, at and below 0, respectively. We then multiplied these values by the -log10 nominal *p*-value. This resulted in a ranked list, where the most strongly upregulated genes were at the top of the list and the most strongly downregulated genes were at the bottom. We used GSEA Preranked (23,24) to conduct the enrichment analysis. Specifically, we tested for enrichment within all Gene Ontology (GO) pathways (i.e., biological processes, cellular component, and molecular function), with at least 5 (and no more than 200) genes within each pathway (8,544 total gene sets). A normalized enrichment score (NES) was calculated for each pathway, and a permutation-based *p*-value (based on 1,000 permutations) was calculated and corrected for multiple testing, resulting in a false-discovery rate (FDR) adjusted *p*-value for each pathway. Statistical significance for enrichment within each pathway was determined by an FDR adjusted *p*-value < .10.

## Results

### Differential Gene Expression

To ascertain differences in gene expression between opioid positive samples compared to controls, we performed RNA-sequencing on postmortem brain specimens from those who died of acute opioid intoxication (n = 72) and group-matched controls (n = 81), as described in the Methods. Differential expression analysis revealed 16 genes that survived multiple test correction (i.e., adj. *p* < .10; see Figure 1), 13 of which were downregulated and three that were upregulated. The top 20 differentially expressed genes are presented in Table 3. Among the top differentially expressed genes, we identified *NPAS4* (log2 fold change = −2.53, *adj. p* = .005), which is implicated in cocaine use (25). There were several other genes that reached or approached significance and are relevant to opioid use. Specifically, we detected two Regulator of G-Protein Signaling genes *(RGS2, adj, p* = .069; *RGS5, adj. p* = .108), one protocadherin gene *(PCDHB12; adj. p* = .108) and an immediate early gene, *EGR4 (adj. p* = .082). Lastly, since the present analysis was able to account for potential key confounders in psychiatric diagnosis and cellular composition, we examined the influence of these covaries on differential expression. There was no evidence of differential gene expression as a result of a psychiatric diagnosis other than a substance use disorder, or by cellular composition *(adj. p’s* > .10; see Figure 2).

**Figure 1.**
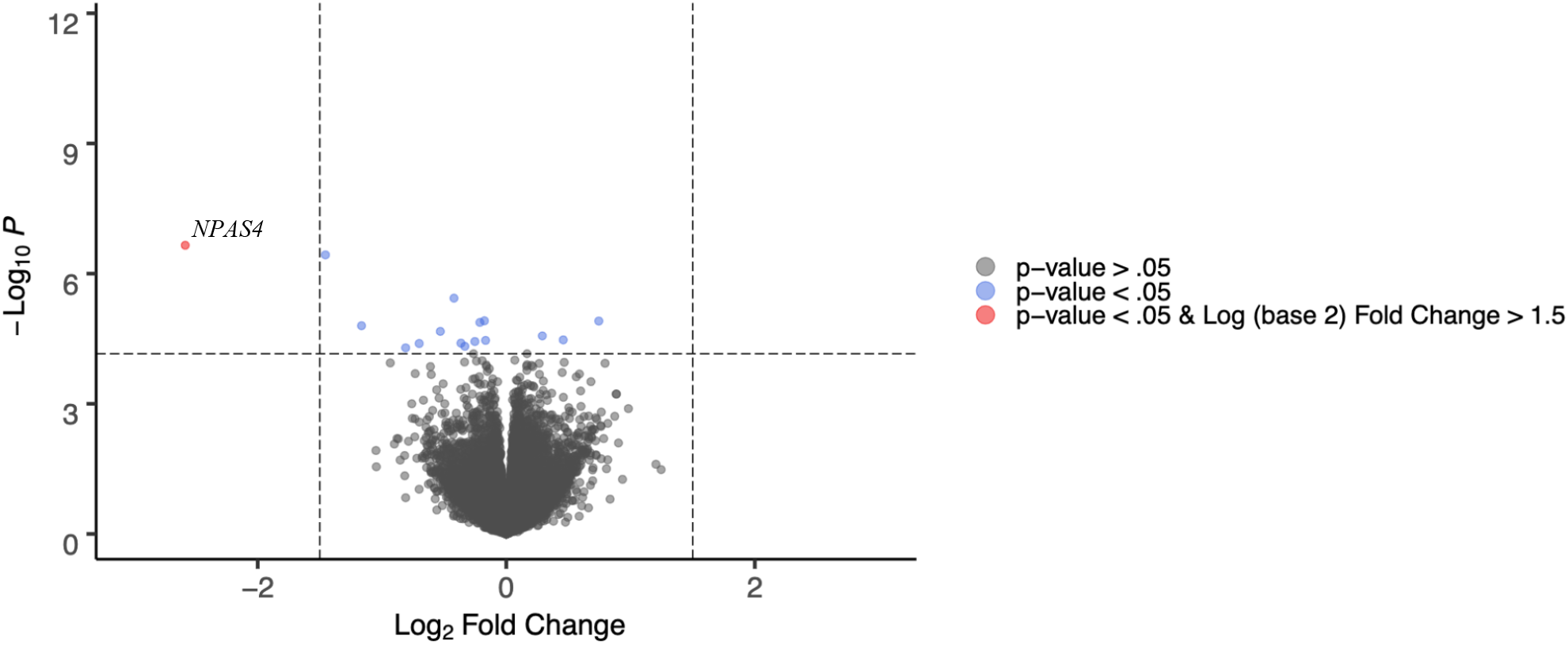
Volcano Plot of Differentially Expressed Genes. *Note.* Horizontal line indicates an FDR-corrected *p*-value of .10 and vertical lines represent a log2 fold change greater or less than +/− 1.5.

**Figure 2.**
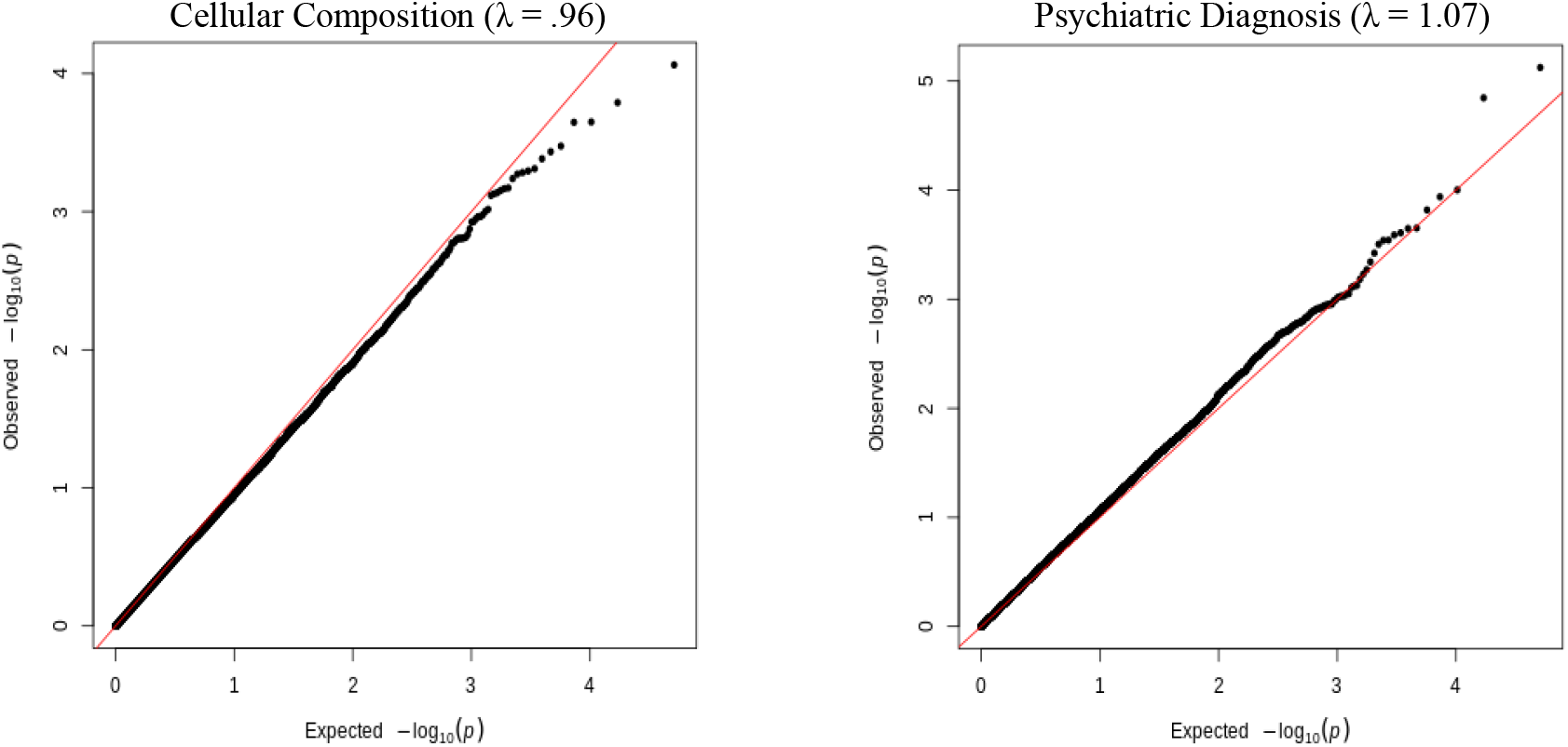
QQ-plots for Psychiatric Diagnosis and Cellular Composition Covariates

**Table 3.** Top 20 Differentially Expressed Genes

| UCSC<br>Gene Symbol | Average<br>Expression | Log Fold-Change | *p*-value | FDR adjusted<br>*p*-value |
| --- | --- | --- | --- | --- |
| *NPAS4* | .340 | -2.583 | 2.22E-07 | .005 |
| *RP11-667K14.3* | -1.184 | -1.455 | 3.70E-07 | .005 |
| *RP11-709D24.8* | .973 | -.421 | 3.68E-06 | .031 |
| *ISG20L2* | 4.027 | -.176 | 1.22E-05 | .057 |
| *ZNF184* | 5.260 | -.212 | 1.33E-05 | .057 |
| *SNORA53* | 2.896 | .744 | 1.24E-05 | .057 |
| *RP11-667K14.4* | -1.652 | -1.165 | 1.59E-05 | .058 |
| *RGS2* | 4.637 | -.530 | 2.15E-05 | .069 |
| *CENPF* | 2.683 | -.365 | 4.01E-05 | .075 |
| *HES1* | 2.900 | .458 | 3.37E-05 | .075 |
| *GPR22* | 5.890 | -.167 | 3.46E-05 | .075 |
| *GATC* | 5.240 | -.253 | 3.67E-05 | .075 |
| *AP000432.1* | -.903 | -.701 | 4.07E-05 | .075 |
| *MT-ND1* | 9.283 | .290 | 2.73E-05 | .075 |
| *RP4-569M23.4* | .629 | -.333 | 4.73E-05 | .081 |
| *EGR4* | 1.122 | -.810 | 5.13E-05 | .082 |
| *PCDHB12* | 3.627 | .167 | 7.01E-05 | .100 |
| *RP11-395G23.3* | 3.270 | -.263 | 7.04E-05 | .100 |
| *RGS5* | 7.268 | .168 | 1.26E-04 | .108 |
| *PTGS2* | 3.821 | -.610 | 1.39E-04 | .108 |

*Note.* Average expression refers to average log<sub>2</sub>-expression for the probe over all arrays and channels.

### Gene Set Enrichment Analysis

Among the most highly expressed genes in the analysis, 10 gene sets were enriched at an FDR corrected *p*-value < .10 (see Table 4). The top GO pathway was related to Bone Morphogenetic Protein (BMP) receptor signaling and no other pathways appeared to be related to opioid use. Among the most strongly downregulated genes in the analysis, 23 gene sets were enriched at an FDR corrected *p*-value < .10 (see Table 5). The top GO pathways were related to protein binding and folding, respectively, and no other pathways appeared to be related to opioid use.

**Table 4.** Significant Results from Gene Set Enrichment Analysis (Upregulated Genes)

| GO ID; Ontology | Term | Adj. *p*-value |
| --- | --- | --- |
| GO:0030510; BP | Regulation of BMP Signaling Pathway | .057 |
| GO:0044224; CC | Juxtaparanode Region of Axon | .067 |
| GO:0098803; CC | Respiratory Chain Complex | .076 |
| GO:0062009; BP | Secondary Palate Development | .087 |
| GO:0030513; BP | Positive Regulation of BMP Signaling Pathway | .088 |
| GO:0045747; BP | Positive Regulation of Notch Signaling Pathway | .089 |
| GO:0003158; BP | Endothelium Development | .093 |
| GO:0045275; CC | Respiratory Chain Complex III | .096 |
| GO:0003222; BP | Ventricular Trabecular Myocardium Morphogenesis | .099 |
| GO:0021889; BP | Olfactory Bulb Interneuron Differentiation | .099 |

*Note.* BP = biological processes; CC = cellular component; MF = molecular function. Significance was determined by an FDR corrected *p*-value < .10.

**Table 5.** Significant Results from Gene Set Enrichment Analysis (Downregulated Genes)

| GO ID; Ontology | Term | FDR *p*-value |
| --- | --- | --- |
| GO:0051082; MF | Unfolded Protein Binding | .002 |
| GO:0006458; BP | De Novo Protein Folding | .002 |
| GO:0061077; BP | Chaperone Mediated Protein Folding | .007 |
| GO:0044183; MF | Protein Folding Chaperone | .007 |
| GO:0042026; BP | Protein Refolding | .017 |
| GO:0042273; BP | Ribosomal Large Subunit Biogenesis | .018 |
| GO:0030684; CC | Pre-ribosome | .018 |
| GO:0051085; BP | Chaperone Cofactor Dependent Protein Refolding | .019 |
| GO:0036499; BP | Perk Mediated Unfolded Protein Response | .019 |
| GO:0006376; BP | mRNA Splice Site Selection | .020 |
| GO:0033549; MF | Map Kinase Phosphatase Activity | .021 |
| GO:0032743; BP | Positive Regulation of Interleukin 2 Production | .023 |
| GO:0030728; BP | Ovulation | .024 |
| GO:0035966; BP | Response to Topologically Incorrect Protein | .027 |
| GO:0000188; BP | Inactivation of MAPK Activity | .029 |
| GO:0101031; CC | Chaperone Complex | .030 |
| GO:003662; MF | Pre mRNA Binding | .040 |
| GO:0045292; BP | mRNA Cis Splicing Via Spliceosome | .044 |
| GO:0051131; BP | Chaperone Mediated Protein Complex Assembly | .046 |
| GO:0051412; BP | Response to Corticosterone | .056 |
| GO:0070202; BP | Regulation of Establishment of Protein Localization to Chromosome | .088 |
| GO:0042255; BP | Ribosome Assembly | .091 |
| GO:0044342; BP | Type B Pancreatic Cell Proliferation | .093 |

*Note.* BP = biological processes; CC = cellular component; MF = molecular function. Significance was determined by an FDR corrected *p*-value < .10.

### Replication of Prior EWAS

In an attempt to validate prior findings, we compared the effect sizes (i.e., log2 fold change) of the top genes identified by Saad et al. (13) to the effect sizes of those same genes in the present analysis. Among the 545 genes differentially expressed in their study, 55 genes were not sequenced in our analysis and 15 did not survive our quality control, leading to 490 genes that overlapped across studies. As can be seen in Figure 3, most differentially expressed genes, including *NPAS4,* were upregulated in Saad et al.’s analysis, whereas most of those genes were slightly downregulated in our analysis. Moreover, it is notable that most of these genes did not reach FDR significance in our study, even though effect sizes did not vary by a large magnitude.

**Figure 3.**
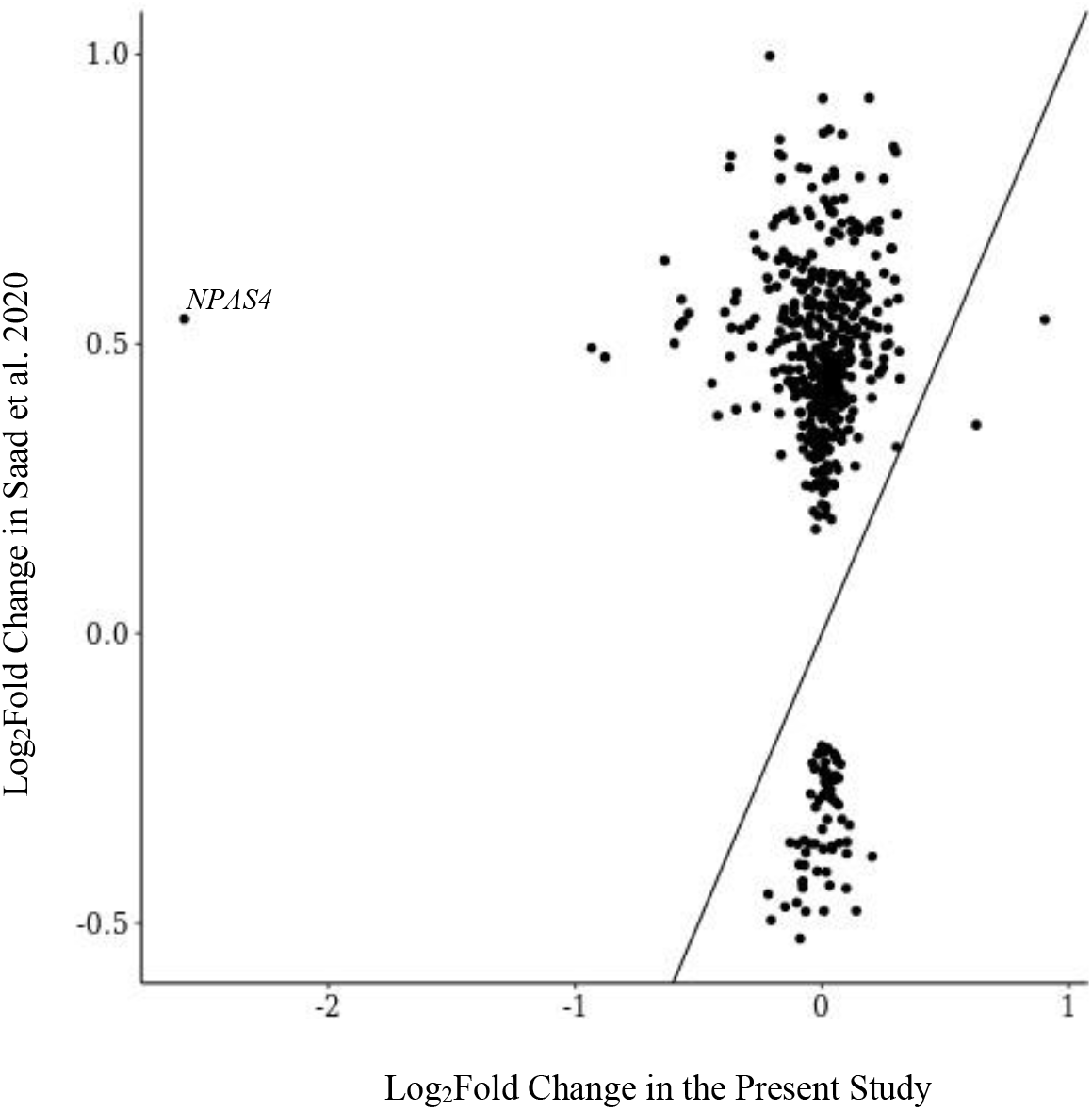
Comparison of Log2 Fold Change in Differentially Expressed Genes Shared with Saad et al. 2020. *Note.* The solid line indicates a slope of 1, meaning that if effect sizes were identical across studies they would fall along the solid line.

## Discussion

The goal of this study was to examine genome-wide differences in gene expression within the human dlPFC following acute opioid intoxication. We detected 16 FDR-significant differentially expressed genes between opioid-positive and control samples. Among these genes was *NPAS4*, which has been identified in preclinical work related to cocaine use (25). Several other genes *(RGS2, RGS5, EGR4)* previously implicated in opioid and cocaine use (26,27) were among the top-ranking genes in our analysis. Gene set enrichment analyses did not reveal enrichment of gene expression within known opioid-relevant gene sets. Nonetheless, the present findings advance our understanding of gene expression affected by opioid overdose and potential targets for therapeutic intervention.

The results support acute opioid intoxication affecting expression of genes directly linked to substance use behaviors. The primary finding from this study was that *NPAS4* was differentially under-expressed in opioid samples compared to controls. Neuronal PAS domain protein 4 *(NPAS4)* is a protein coding gene that influences expression of brain-derived neurotrophic factor (*BNDF*) (28). It has been shown to control GABAnergic synapse development in an activity-dependent manner, and GABAergic deficits have been implicated in psychiatric disorders such as substance use disorders and major depressive disorder (29). Recent work examining the NAc of Sprague Dawley rats showed that cocaine inhibits *HDAC5* – a histone deacetylase enzyme – from entering the cell nucleus and limits production of *Npas4;* subsequent up-regulation of *Npas4* is associated with increased drug-environment associations (25). Interestingly, *NPAS4* was downregulated in the present analysis. Preclinical evidence suggests that acute opioid use is associated with upregulation of *Npas4* (27,30). However, preclinical studies of methamphetamine use suggest that *Npas4* is significantly downregulated hours after initial upregulation (31). The reasons for the discrepancy in findings related to *NPAS4* make interpretation of this finding difficult. It is possible that the opioid users in the present sample were chronic users, but psychiatric histories obtained for the samples only indicated lifetime diagnosis of psychiatric disorders.

It also is interesting to note several other genes implicated in substance use that were significant (i.e., FDR adjusted *p* ≤ .10). Specifically, *PCDHB12, RGS2,* and *EGR4* were among the top differentially expressed genes in our analysis. Protocadherin Beta 12 *(PCHDB12’)* is a gene within the protocadherin gene cluster; this gene cluster is responsible for intracellular signaling and cell adhesion. Prior work examining nicotine withdrawal in humans discovered that a variant (rs31746) within the PCDH-α, -β, -γ gene cluster was associated with nicotine cravings and likelihood of relapse based on self-reported smoking behavior. Furthermore, this variant was associated with differential expression of the *PCDHβ8* gene in postmortem PFC tissue of humans (32). Although there is no other evidence of differential expression of protocadherin genes in relation to opioid use, the present findings suggest they may play a role in acute opioid intoxication.

The detection of differential expression among regulator of G protein signaling 2 and 5 (*RGS2*, *RGS5*) genes are particularly relevant given their known G-protein coupled receptors, namely μ-opioid receptors (33). Sutton et al. (33) demonstrated that, in response to morphine administration, deletion of *Rgs7* increased reward and analgesia, while heightening withdrawal, in mice. Further exploration of *RGS2* and *RGS5* is warranted as these genes may be reasonable targets for medications to treat OUD. Lastly, detection of differential expression in early growth response gene 4 (*EGR4*) is interesting given the role of immediate early genes. Immediate early genes are responsible for encoding transcription factors, which can interact with gene promoters to effect changes in gene expression at downstream genes (27). Preclinical evidence suggests that *Egr1*, which is in the same gene family at *EGR4*, is differentially expressed in the forebrain of mice after morphine administration (34). Further examination of these genes and their potential role in opioid abuse is warranted and may elucidate alternative mechanisms through which opioid dependence, cravings, and other drug-related behaviors develop and persist.

### Strengths and Limitations

The primary strength of this study is the assessment of gene expression in the dlPFC of human subjects. To date, only one other published study (13) has assessed gene expression in brains of human opioid users, focusing on the midbrain. Although the top 100 differentially expressed genes did not overlap with those in Saad et al.’s study, our samples reflect gene expression following an overdose (as opposed to chronic use), which may explain some of the lack of concordance in differentially expressed genes between the two studies. Nonetheless, these findings shed light on patterns of gene expression in a different brain region implicated in opioid use disorders. Additional work with postmortem samples will clarify similarities and differences in gene expression across brain regions implicated in addiction. Another strength of this study is the use of transcriptome-wide analysis of gene expression. This allows for a more complete view of the influence of opioid use on gene expression outside of candidate genes such as μ-opioid receptors. Lastly, the study benefitted from the inclusion of cellular composition and a sample of group-matched psychiatric controls. Although the current findings do not provide evidence for their direct influence on gene expression, inclusion of these variables accounts for key confounders that allows for a more accurate analysis and interpretation of the association between opioid use and gene expression.

This study also has several limitations to consider when interpreting the findings. First, although opioid users died of acute opioid intoxication and were determined to have a lifetime diagnosis of OUD, the course of opioid misuse among those individuals remains unclear (e.g., age of onset, frequency of use). This presents a challenge in teasing out effects due to chronic versus acute opioid misuse. Examination of tissue within a single brain region also present a minor limitation in interpretation of results. Although the PFC is implicated in addiction, the NAc is often the focus in studies of addiction (5,6). Given the link between the PFC and NAc in addiction, it will be important to replicate the current study across various brain regions. Lastly, future studies will benefit from more racially and ethnically diverse samples, as well as samples including individuals across various developmental stages. The latter, in particular, may elucidate differential effects of opioid abuse depending on brain maturation and aging.

### Conclusions

This study represents the first genome-wide analysis of gene expression in the dlPFC of opioid users and group-matched controls. Evidence suggests that opioid use is associated with differential expression of several genes, including *NPAS4,* which is implicated in drug cravings and drug-environment cues. Additional genes implicated in substance abuse provide promising avenues for continued research. Future studies examining chronic versus acute opioid use and polysubstance use, and those with larger, more racially and developmentally diverse samples will clarify these associations and aid in determining the value of differentially expressed genes as therapeutic targets for substance use disorders.

## Acknowledgements

The authors would like to express their gratitude to our colleagues whose efforts have led to the donation of postmortem tissue to advance these studies, including at the Office of the Chief Medical Examiner of the State of Maryland, Baltimore Maryland and the Office of the Chief Medical Examiner of Kalamazoo County Michigan. We also would like to acknowledge the contributions of Dr. Llewelyn Bigelow for his diagnostic expertise. Finally, we are indebted to the generosity of the families of the decedents, who donated the brain tissue used in these studies.

## References

1. O’donnell J, Gladden; R Matt, Mattson CL, Hunter CT, Davis NL. Morbidity and Mortality Weekly Report Vital Signs: Characteristics of Drug Overdose Deaths Involving Opioids and Stimulants-24 States and the District of Columbia [Internet]. 2019 [cited 2020 Sep 17]. Available from: https://www.cdc.gov/drugoverdose/od2a/index.html.

2. Scholl L, Seth P, Kariisa M, Wilson N, Baldwin G. Morbidity and Mortality Weekly Report Drug and Opioid-Involved Overdose Deaths-United States [Internet]. Vol. 67, MMWR Morbidity and Mortality Weekly Report. 2019 [cited 2020 Sep 14]. Available from: https://www.cdc.gov/nchs/data/nvsr/nvsr61/

3. Florence CS, Zhou C, Luo F, Xu L. The economic burden of prescription opioid overdose, abuse, and dependence in the United States, 2013. Med Care. 2016;54(10):901–6.

4. Egervari G, Kozlenkov A, Dracheva S, Hurd YL. Molecular windows into the human brain for psychiatric disorders [Internet]. Vol. 24, Molecular Psychiatry. 2019 [cited 2020 Sep 17]. p. 653–73. Available from: https://doi.org/10.1038/s41380-018-0125-2

5. Volkow ND, Morales M. The Brain on Drugs: From Reward to Addiction [Internet]. Vol. 162, Cell. Cell Press; 2015 [cited 2020 Sep 17]. p. 712–25. Available from: http://dx.doi.org/10.1016/j.cell.2015.07.046

6. Kalivas PW, Volkow N, Seamans J. Unmanageable motivation in addiction: A pathology in prefrontal-accumbens glutamate transmission. Vol. 45, Neuron. Cell Press; 2005. p. 647–50.

7. Kruyer A, Chioma VC, Kalivas PW. The Opioid-Addicted Tetrapartite Synapse. Vol. 87, Biological Psychiatry. Elsevier USA; 2020. p. 34–43.

8. Johnson SW, North RA. Opioids excite dopamine neurons by hyperpolarization of local interneurons. J Neurosci. 1992;12(2):483–8.

9. Browne CJ, Godino A, Salery M, Nestler EJ. Epigenetic Mechanisms of Opioid Addiction. Vol. 87, Biological Psychiatry. Elsevier USA; 2020. p. 22–33.

10. Jones JD, Vadhan NP, Luba RR, Comer SD. The effects of heroin administration and drug cues on impulsivity. J Clin Exp Neuropsychol [Internet]. 2016 Jul 2 [cited 2020 Sep 17];38(6):709–20. Available from: https://www.tandfonline.com/doi/full/10.1080/13803395.2016.1156652

11. Arias F, Arnsten JH, Cunningham CO, Coulehan K, Batchelder A, Brisbane M, et al. Neurocognitive, psychiatric, and substance use characteristics in opioid dependent adults. Addict Behav. 2016 Sep 1;60:137–43.

12. Biernacki K, McLennan SN, Terrett G, Labuschagne I, Rendell PG. Decision-making ability in current and past users of opiates: A meta-analysis. Vol. 71, Neuroscience and Biobehavioral Reviews. Elsevier Ltd; 2016. p. 342–51.

13. Saad MH, Rumschlag M, Guerra MH, Savonen CL, Jaster AM, Olson PD, et al. Differentially expressed gene networks, biomarkers, long noncoding RNAs, and shared responses with cocaine identified in the midbrains of human opioid abusers. Sci Rep [Internet]. 2019 Dec 1 [cited 2020 Sep 17];9(1):1–9. Available from: https://doi.org/10.1038/s41598-018-38209-8

14. Kang HJ, Kawasawa YI, Cheng F, Zhu Y, Xu X, Li M, et al. Spatio-temporal transcriptome of the human brain. Nature [Internet]. 2011 Oct 27 [cited 2020 Sep 17];478(7370):483–9. Available from: https://www.nature.com/articles/nature10523

15. Collado-Torres L, Burke EE, Peterson A, Shin JH, Straub RE, Rajpurohit A, et al. Regional Heterogeneity in Gene Expression, Regulation, and Coherence in the Frontal Cortex and Hippocampus across Development and Schizophrenia. Neuron [Internet]. 2019 Jul 17 [cited 2020 Nov 12];103(2):203–216.e8. Available from: /pmc/articles/PMC7000204/?report=abstract

16. Kim D, Paggi JM, Park C, Bennett C, Salzberg SL. Graph-based genome alignment and genotyping with HISAT2 and HISAT-genotype. Nat Biotechnol [Internet]. 2019 Aug 1 [cited 2020 Nov 12];37(8):907–15. Available from: https://pubmed.ncbi.nlm.nih.gov/31375807/

17. Liao Y, Smyth GK, Shi W. FeatureCounts: An efficient general purpose program for assigning sequence reads to genomic features. Bioinformatics [Internet]. 2014 Apr 1 [cited 2020 Nov 12];30(7):923–30. Available from: https://pubmed.ncbi.nlm.nih.gov/24227677/

18. Team RC. R: A language and environment for statistical computing [Internet]. Vienna: R Foundation for Statistical Computing; 2018. Available from: https://www.r-project.org/

19. Ritchie ME, Phipson B, Wu D, Hu Y, Law CW, Shi W, et al. Limma powers differential expression analyses for RNA-sequencing and microarray studies. Nucleic Acids Res [Internet]. 2015 Jan 6 [cited 2020 Sep 16];43(7):e47. Available from: https://academic.oup.com/nar/article/43/7/e47/2414268

20. Jaffe AE, Tao R, Norris AL, Kealhofer M, Nellore A, Shin JH, et al. QSVA framework for RNA quality correction in differential expression analysis. Proc Natl Acad Sci U S A [Internet]. 2017 Jul 3 [cited 2020 Sep 21];114(27):7130–5. Available from: https://www.pnas.org/content/114/27/7130

21. Benjamini Y, Hochberg Y. Controlling the False Discovery Rate: A Practical and Powerful Approach to Multiple Testing. J R Stat Soc Ser B [Internet]. 1995 Jan 1 [cited 2020 Sep 18];57(1):289–300. Available from: https://rss.onlinelibrary.wiley.com/doi/full/10.1111/j.2517-6161.1995.tb02031.x

22. Reimand J, Isser R, Voisin V, Kucera M, Tannus-Lopes C, Rostamianfar A, et al. Pathway enrichment analysis and visualization of omics data using g:Profiler, GSEA, Cytoscape and EnrichmentMap. Nat Protoc. 2019;14(2):482–517.

23. Mootha VK, Lindgren CM, Eriksson KF, Subramanian A, Sihag S, Lehar J, et al. PGC-lα-responsive genes involved in oxidative phosphorylation are coordinately downregulated in human diabetes. Nat Genet [Internet]. 2003 Jul 1 [cited 2020 Oct 17];34(3):267–73. Available from: https://www.nature.com/articles/ng1180

24. Subramanian A, Tamayo P, Mootha VK, Mukherjee S, Ebert BL, Gillette MA, et al. Gene set enrichment analysis: A knowledge-based approach for interpreting genome-wide expression profiles. Proc Natl Acad Sci U S A [Internet]. 2005 Oct 25 [cited 2020 Oct 17];102(43):15545–50. Available from: www.pnas.orgcgidoi10.1073pnas.0506580102

25. Taniguchi M, Carreira MB, Cooper YA, Bobadilla AC, Heinsbroek JA, Koike N, et al. HDAC5 and Its Target Gene, Npas4, Function in the Nucleus Accumbens to Regulate Cocaine-Conditioned Behaviors. Neuron [Internet]. 2017 Sep 27 [cited 2020 Sep 18]; 96(1):130–144.e6. Available from: https://pubmed.ncbi.nlm.nih.gov/28957664/

26. Senese NB, Kandasamy R, Kochan KE, Traynor JR. Regulator of G-Protein Signaling (RGS) Protein Modulation of Opioid Receptor Signaling as a Potential Target for Pain Management [Internet]. Vol. 13, Frontiers in Molecular Neuroscience. Frontiers Media S.A.; 2020 [cited 2020 Oct 17]. p. 5. Available from: www.frontiersin.org

27. Bisagno V, Cadet JL. Expression of immediate early genes in brain reward circuitries: Differential regulation by psychostimulant and opioid drugs [Internet]. Vol. 124, Neurochemistry International. 2019 [cited 2020 Sep 23]. p. 10–8. Available from: https://doi.org/10.1016/j.neuint.2018.12.004

28. Lin Y, Bloodgood BL, Hauser JL, Lapan AD, Koon AC, Kim TK, et al. Activity-dependent regulation of inhibitory synapse development by Npas4. Nature [Internet]. 2008 [cited 2020 Sep 23];455(7217):1198–204. Available from: https://idp.nature.com/authorize/casa?redirect_uri= https://www.nature.com/articles/nature_07319&casa_token=aGqkFSABU_EAAAAA:-4HEhUSVMFOtbg0omzXc_kVaupp0a1PYGiwY01SoK44TuPhugRJ1cJDLrDRFJl7dcIB_xp8scEnTZ4kSI

29. Luscher B, Shen Q, Sahir N. The GABAergic deficit hypothesis of major depressive disorder [Internet]. Vol. 16, Molecular Psychiatry. 2011 [cited 2020 Sep 23]. p. 383–406. Available from: https://www.nature.com/articles/mp2010120

30. Piechota M, Korostynski M, Solecki W, Gieryk A, Slezak M, Bilecki W, et al. The dissection of transcriptional modules regulated by various drugs of abuse in the mouse striatum. Genome Biol [Internet]. 2010 May 4 [cited 2020 Sep 23];11(5):48. Available from: http://genomebiology.com/2010/11/5/R48

31. Martin TA, Jayanthi S, McCoy MT, Brannock C, Ladenheim B, Garrett T, et al. Methamphetamine causes differential alterations in gene expression and patterns of histone acetylation/hypoacetylation in the rat nucleus accumbens. Manzoni OJ, editor. PLoS One [Internet]. 2012 Mar 28 [cited 2020 Sep 23];7(3):e34236. Available from: https://dx.plos.org/10.1371/journal.pone.0034236

32. Jensen KP, Smith AH, Herman AI, Farrer LA, Kranzler HR, Sofuoglu M, et al. A protocadherin gene cluster regulatory variant is associated with nicotine withdrawal and the urge to smoke. Mol Psychiatry [Internet]. 2017 Feb 1 [cited 2020 Sep 24];22(2):242–9. Available from: http://locuszoom.sph.

33. Sutton LP, Ostrovskaya O, Dao M, Xie K, Orlandi C, Smith R, et al. Regulator of G-Protein Signaling 7 Regulates Reward Behavior by Controlling Opioid Signaling in the Striatum. Biol Psychiatry. 2016 Aug 1;80(3):235–45.

34. ZióŁkowska B, Gieryk A, Solecki W, PrzewŁocki R. Temporal and anatomic patterns of immediate-early gene expression in the forebrain of C57BL/6 and DBA/2 mice after morphine administration. Neuroscience. 2015 Jan 2;284:107–24.

